# Molecular Characterization and Antibiotic Resistance Profile of *Escherichia coli* Isolated From Liver Abscess

**DOI:** 10.1101/2024.05.23.595549

**Authors:** Mohammad Nasar, Sarrar Grazza

**Author notes:** Correspondence Author: Mohammad Nasar.

## Abstract

**Background:** Bacterial liver abscess is the most common hepatic infection, which can lead to death. *Escherichia coli* is among the many species of bacteria that cause it. This study was conducted to isolate *E. coli* from liver abscess and then to characterise the bacteria’s molecular makeup and antibiotic resistance profile.

**Methods:** A total of 208 stool samples were collected from patients showing symptoms of liver abscess. *E. coli* was isolated from these samples followed by identification by biochemical tests. Pure and biochemically positive colonies were confirmed by polymerase chain reaction. The disk diffusion method was used to ascertain the pattern of antibiotic resistance exhibited by *E. coli* isolates.

**Results:** The PCR amplification efficiency was nearly 100% since all of the samples appeared at 284 molecular base pairs (bp), which is considered to be the optimal parameter assay. The antimicrobial susceptibility pattern showed that isolates were resistant to many drugs but 100% and 92% of the isolates were susceptible to imipenem and azithromycin, respectively. All isolates were resistant to ampicillin, vancomycin, and cefotaxime. This was followed by ceftazidime (72%), tetracycline (84%), trimethoprim (80%), streptomycin (96%), linezolid (92%), Teicoplanin (80%), nalidixic acid (84%), ciprofloxacin (92%), and chloramphenicol (72%).

**Conclusion:** Multiple drug resistant *E. coli* is one of the causes of liver abscesses in humans.

## 1. Introduction

Enterobacteriales, a bacterial order in the phylum Proteobacteria, consists of several important intestinal pathogens (Bujňáková et al., 2022). This order includes persistent gut colonizers that make up small microbiota components under healthy conditions (Riccio and Rossano, 2020). As long as the microbiota is in balance and the complex and dense bacterial population inhibits their overgrowth, opportunistic Enterobacteria can continue to exist as gut commensals without causing any infections (Rao et al., 2020). A disruption of the microbiota can result in a bloom of Enterobacteria, which can cause pathogen-mediated illnesses and inflammatory reactions in the host (Amaretti et al., 2020).

The most studied species among the Enterobacteria is *Escherichia coli* in terms of the traits that differentiate pathogenicity and commensalism (Dalmasso et al., 2023). It mostly colonizes the intestine, but it also includes many pathogenic variations that can cause infections in other tissues or the gut, in addition to harmless commensals. *E. coli* is also found in patients with liver complications such as ascites or urinary tract infections. Patients with liver disease have a decreased ability to fight against bacterial infections, which exposes them to risk of infections, sepsis, and even death. Among these patients, spontaneous bacterial peritonitis, bacteremia, skin and soft tissue infections, pneumonia and urinary tract infections are the most frequent bacterial illnesses (Bunchorntavakul et al., 2016). The most common causes are Gram negative bacteria. The majority of research interest has been focused on virulent strains of *E. coli* isolated from infected patients (Dalmasso et al., 2023), but there has also been a growing focus on environmental strains and faecal isolates from healthy subjects in an effort to assess the pathogenic potential of a larger reservoir of biodiversity (Keesing and Ostfeld, 2021).

*Escherichia coli* is a gram-negative, rod-shaped, facultatively anaerobic bacterium, which is usually found in the lower intestine of worm-blooded species that are endotherms (Tenaillon et al., 2010). Living in the lower digestive tract, *E. coli* is present within the first 24 hours of birth (Bettelheim and Lennox-King, 1976). The gut microbiota is made up of over 500 different species of bacteria, with 10^10^–10^11^ cells per gram of large-intestinal content. Along with other facultative anaerobes, *E. coli* makes up around 0.1% of the local gut flora (Berg, 1996; Eckburg et al., 2005). As a normal component of the flora, it is primarily regarded as helpful to humans (Croxen and Finlay, 2010). *E. coli* has the unusual ability to be utilized as an indicator organism to check for microbial contamination in water sources (Devane et al., 2020). Certain pathogenic strains are also present. The pathogenicity of a given pathotype is determined by the presence of a set of virulence factors, which facilitate the infection of humans and animals with the bacteria and the manifestation of specific symptoms (Watkins et al., 2016).

Antimicrobials are usually used for treating infected patients and also for prophylaxis for certain ailments. The antimicrobial abuse and inadequate selection are the key reasons for the emergence of resistance among several bacteria and this makes the antibacterial therapy more difficult (FR et al., 2019). Since *E. coli* are commensal bacteria, they are thought to be a reservoir of pathogenic bacteria’s resistance genes (Lambrecht et al., 2019). Their degree of resistance is thought to be a useful indicator of the selection pressure brought on by the administration of antibiotics as well as the likelihood that these pathogens may have resistance challenges (Hoang et al., 2017). In addition to transmit antibiotic resistance genes to other *E. coli* strains, resistant strains of the bacterium can also pick up resistance from other species and transfer it to other bacteria in the gastrointestinal tract (Tawfick et al., 2022). Thus, determining the prevalence of *E. coli* in liver abscess as well as characterizing the bacteria’s molecular makeup and antibiotic resistance profile was the main objective of this study.

## 2. Materials and Methods

### 2.1. Culture media and chemicals

Luria broth, nutrient agar, beef extract, MacConkey agar, Simmon’s citrate agar, crystal violet, and antibiotic discs (ampicillin, tetracycline, streptomycin, cephotaxime, azithromycin, chloramphenicol, linezolid, teicoplanin, nalidixic acid, ciprofloxacin, trimethoprim and imipenem)., St Louis, MO, USA. Sodium chloride, were purchased from Sigma Chemical Co, hydrogen peroxide, dextrose, lactose, and glucose were procured from Merck, Darmstadt, Germany. All other chemicals were from Shandong Chemicals, China and were of the highest grade available.

### 2.2. Sample collection and Enrichment

Early morning stool samples (∼3 g) were collected from individuals at their residence in sterile blue cap stool containers. The sample was enriched by adding fecal sample (20μl) to 2ml of Luria broth followed by incubation overnight at 37°C.

### 2.3. Isolation and purification

The samples were initially grown (18–24 hours) in nutrient broth at 37ºC and then they were sub-cultured using the streak plate method onto MacConkey agar (Zinnah et al., 2007). The culture was incubated at 37°C for 24 hours until the pure culture with homogenous colonies were obtained. Isolated red/pink colonies were re-streaked on the same agar. The re-streaked petri plates were incubated for 24 hours at 37°C for further purification.

### 2.4. Identification

The purified colonies on re-streaked MacConkey agar were further identified by Gram staining (Kohlerschmidt et al., 2021) and by different biochemical tests.

#### 2.4.1. Biochemical Tests

Bacterial strains were biochemically identified by various tests, e.g., catalase (Chauhan et al., 2020), oxidase (Horne et al., 2024), indole (Alves et al., 2006), methyl red (Shoaib et al., 2020), citrate (Rahman et al., 2021), triple sugar iron agar (Zinnah et al., 2007), and Voges-Proskauer (Zinnah et al., 2007).

### 2.5. DNA Extraction

After 24h incubation the broth was centrifuged at 12,000g for 5 min. The supernatant was discarded and distilled water was added to the pellet. After 5 sec vortexing, the samples were transferred to Eppendorf tube followed by centrifugation at 12,000 rpm for 5 min. The pellet was washed and centrifuged again. After removing supernatant, the pellet was resuspended in 300μl 10% Chelex. After 30 min incubation at 99°C, supernatant was isolated by centrifugation at 12000rpm for 5min (Mahmoud et al., 2020).

### 2.6. PCR amplification

Pure and biochemically positive colonies were confirmed by polymerase chain reaction. Using the Sul 1 primer pairs forward (CGCACCGGAAACATCGCTGCAC) and reverse (TGAAGTTCCGCCGCAAGGCTCG), resistant gene segments in bacterial positive isolates (∼284 bp) were amplified using PCR. The PCR mixes were prepared according to manufacturer’s directions. The PCR amplification was conducted in a 25.0 μl reaction mixture, consisting of 12.5 μl of master mix, 2.0 μl of DNA, 1.5 μl of each primer and 7.5 μl of distilled water. Thermal cycling machine was set up to amplify DNA for 35 cycles. The PCR was initially denaturated for 15 minutes at 95°C. Next, it was denaturated and annealed for 1 minute at 94°C and 52°C, respectively. Finally, it was extended for 1 minute at 72°C and concluded the PCR with a final extension for 15 minutes at 72°C (Mahmoud et al., 2020).

### 2.7. Gel Electrophoresis

PCR was followed by gel electrophoresis to visualize the bands. Extracted DNA (5 μl) was added to each well after adding ethidium bromide (1.5 μl) into agarose gel (2%) and pouring into the casting tray to solidify. The current was adjusted to 100 amps for 30 min while the applied voltage was 60 volts. Following that, a UV transilluminator was used to view the PCR result (Mahmoud et al., 2020).

### 2.8. Antibiotic Susceptibility Test

The Kirby-Bauer disc diffusion method was used to screen the 208 isolates for antibiotic resistance. Fourteen antibiotics were tested, e.g., ampicillin, vancomycin, cefotaxime, ceftazidime, imipenem, tetracycline, trimethoprim, streptomycin, linezolid, Teicoplanin, nalidixic acid, ciprofloxacin, chloramphenicol, and azithromycin. The recommendations’ stated antibiotic minimum inhibitory concentration (MIC) was used to classify *E. coli* patterns as “resistant”, “intermediate resistant,” or “sensitive”. The word “resistant” was applied to all isolates exhibiting “resistant” or “intermediate resistant” patterns (Mahmoud et al., 2020).

## 3. Results

### 3.1. Isolation of *E. coli*

Two hundred eight samples were taken and successfully grown on MacConkey agar media plus Cefotaxime. Ninety one samples showed positive results which exhibit 43% samples were identified as *E. coli*.

### 3.2. Biochemical tests

The biochemical characteristics of E. coli strains are shown in Table 4. In biochemical study all the isolates revealed positive reaction in catalase, oxidase and TSI, with the production of acid and gas within 24-48 hrs of incubation. The isolates also showed negative reaction in Simmon’s citrate and VP test and differential results in Indole and methyl red test.

### 3.3. Polymerase chain reaction (PCR)

The PCR results showed that all subject’s DNA was visible at the 284 bp DNA molecular ladder. According to the findings, DNA was amplified to 284 bp and aligned using a DNA molecular ladder (Figure 6).

**Figure 3.**
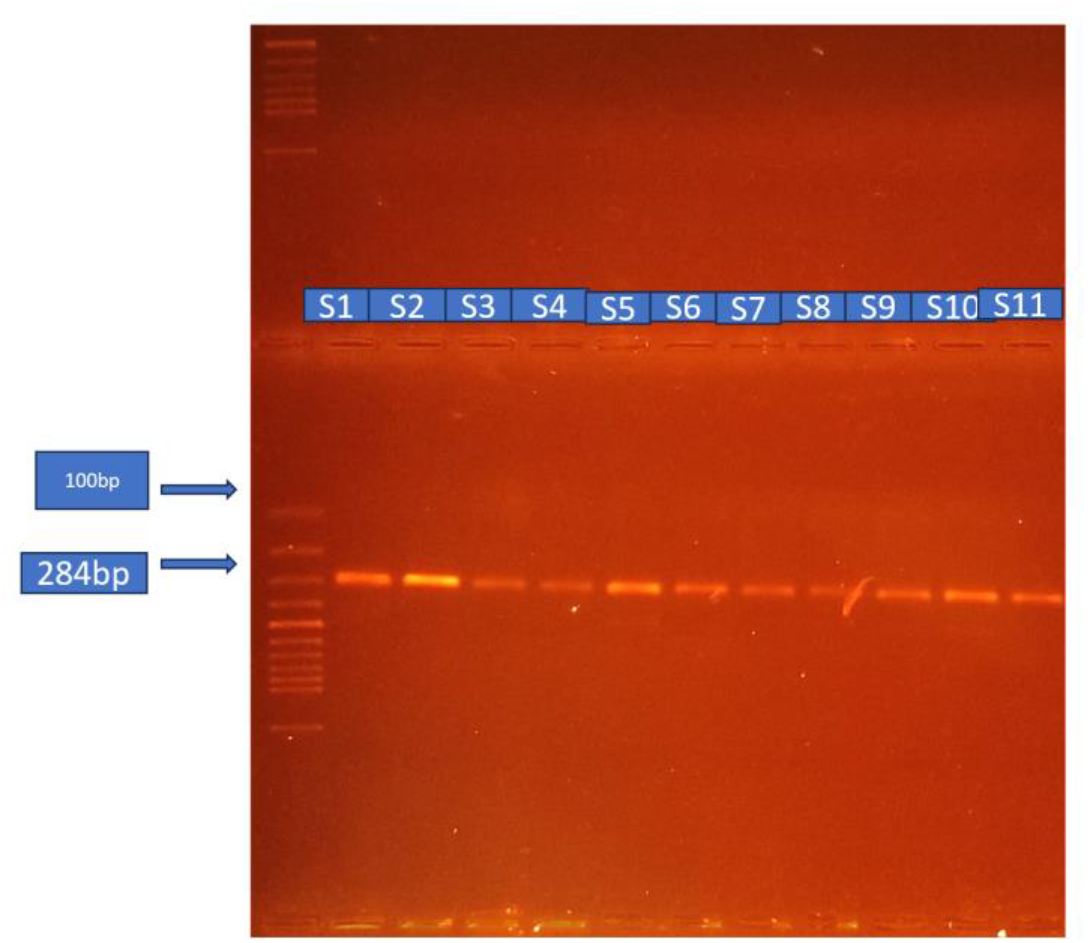
PCR results after DNA molecular ladder analysis and agarose gel electrophoresis.

### 3.4. Phenotypic confirmation of extended spectrum beta-lactamase (ESBL) *E. coli*

The zone of inhibition of corner antibiotics (Aug and TZP) diffused into the center antibiotics (CRO) showed positive results for ESBL production (20-25 mm from corner to center) as shown in figure 4. Table 5 shows antimicrobial susceptibility pattern of *E. coli*. The antimicrobial susceptibility pattern showed that isolates were resistant to most of the drugs, however 100% of isolates were susceptible to imipenem and 92% to azithromycin respectively. The isolates showed resistance to ampicillin (100%), vancomycin (100%) and cefotaxime (100%), were the highest among the antibiotics used, followed by ceftazidime (72%), tetracycline (84%), trimethoprim (80%), streptomycin (96%), linezolid (92%), Teicoplanin (80%), nalidixic acid (84%), ciprofloxacin(92%) and chloramphenicol (72%).

**Table 5.** Antimicrobial susceptibility pattern of *E. coli*.

| S. No. | Antibiotics | Resistance (%) | Sensitivity (%) | Intermediate (%) |
| --- | --- | --- | --- | --- |
| 1. | Ampicillin | 100 | 0 | 0 |
| 2. | Cefotaxime | 100 | 0 | 0 |
| 3. | Ceftazidime | 72 | 28 | 0 |
| 4. | Tetracycline | 84 | 16 | 0 |
| 5. | Trimethoprim | 80 | 20 | 0 |
| 6. | Imipenem | 0 | 100 | 0 |
| 7. | Chloramphenicol | 72 | 24 | 4 |
| 8. | Streptomycin | 96 | 0 | 4 |
| 9. | Linezolid | 92 | 0 | 8 |
| 10. | Teicoplanin | 80 | 0 | 20 |
| 11. | Vancomycin | 100 | 0 | 0 |
| 12. | Nalidixic Acid | 84 | 0 | 16 |
| 13. | Ciprofloxacin | 92 | 0 | 8 |
| 14. | Azithromycin | 4 | 92 | 4 |

## 4. Discussion

Multidrug resistant *E. coli* has grown to be a concerning issue and has been observed in increasing numbers in humans. *E. coli* is a common cause of contaminated drinking water, which can result in significant complications such liver abscess, diarrhoea, and enteritis. It is also one of the main etiologic agents causing urinary tract infections, sepsis, enteritis, and meningitis (Su and Brandt, 1995). Global public health is greatly affected by antibiotic resistant *E. coli* (Galindo-Méndez, 2020; Puvača and de Llanos Frutos, 2021). These resistant bacteria could raise the risk to human health (Nji et al., 2021; Ramos et al., 2020). Therefore, it is critical to get further knowledge about such issues.

The antimicrobial susceptibility pattern of the isolates in this investigation revealed that they were resistant to a majority of drugs tested; however, 100% of the isolates were responsive to imipenem and 92% to azithromycin. Among the antibiotics used, the isolates exhibited the highest levels of resistance to ampicillin (100%), vancomycin (100%), and cefotaxime (100%). These were followed by ceftazidime (72%), tetracycline (84%), trimethoprim (80%), streptomycin (96%), linezolid (92%), Teicoplanin (80%), nalidixic acid (84%), ciprofloxacin (92%), and chloramphenicol (72%). These bacterial species can acquire a large number of resistance genes, primarily by horizontal gene transfer (Zarei-Baygi and Smith, 2021).

The Gram stain is a commonly utilized technique for the identification and differentiation of bacteria (Budin et al., 2012). This method is widely applied in clinical diagnostics, environmental sample detection, and identification of bacterial species. Since crystal violet binds to both Gram-positive and Gram-negative bacteria’s peptidoglycan layer, it is employed in the staining procedure for bacterial samples. Crystal violet produces an insoluble complex when it comes into contact with an iodine solution (Becerra et al., 2016; Biswas et al., 1970; Claus, 1992). A thick peptidoglycan coating covers Gram-positive bacteria, while lipopolysaccharides and lipoproteins cover Gram-negative bacteria. Gram-negative bacteria lose colour when decolorized with acetone or alcohol, Gram-positive bacteria retain their purple colour (Kohlerschmidt et al., 2021). The isolates of bacteria were Gram negative based on our Gram staining results. Staining is a reliable and simple procedure, but the final detection is still done by optical microscopy, which is often susceptible to user-dependent sampling error.

In this study, *E. coli* were isolated and identified through conventional microbiological analysis. The outcomes of the microbiological detection of *E. coli* were comparable since, in addition to culture-based detection, the selected isolates’ molecular identity was determined by PCR amplification using 16s rDNA. To establish specificity, we optimized primer and annealing temperature for DNA amplification using a standard set of samples and precise calculations based on nucleotide presence in the study’s DNA sequence. The gradient temperature in this experiment was determined using the forward and reverse primers.

The PCR amplification efficiency was very nearly 100% since all of the samples appeared at 284 molecular base pairs (bp), which is considered to be the ideal parameter assay. A pilot study would require an amplification efficiency of 90% – 105%. Inadequate primer design or less-than-ideal reaction conditions in relation to the PCR’s components can lead to low reaction efficiency. (Bunu et al., 2020). Agarose gel electrophoresis of all the amplified products produced bands that were positive for identification. All of the samples were confirmed to be E. coli using molecular identification.

## Conclusions

Antimicrobial susceptibility assay revealed that the most prevalent resistance patterns against ampicillin, vancomycin, tetracycline, cefotaxime, streptomycin, linezolid, and ciprofloxacin. These were followed by nalidixic acid, Teicoplanin, trimethoprim, and chloramphenicol. On the other hand, azithromycin and imipenem were the most effective antimicrobials. It can be concluded that *E. coli* showed anti-biotic resistance in liver abscess.

## Ethical approval

Ethical approval was not applicable.

## Acknowledgement

The authors thank the Department of Biosciences COMSATS University Islamabad for all the support provided.

